# Apical and basal auxin sources pattern shoot branching in a moss

**DOI:** 10.1101/2022.01.04.474977

**Authors:** Mattias Thelander, Katarina Landberg, Arthur Muller, Gladys Cloarec, Nik Cunniffe, Stéphanie Huguet, Ludivine Soubigou-Taconnat, Véronique Brunaud, Yoan Coudert

## Abstract

Shoot branching mechanisms where branches arise in association with leaves – referred to as lateral or axillary branching – evolved by convergence in the sporophyte of vascular plants and the gametophyte of bryophytes, and accompanied independent events of plant architectural diversification^1^. Previously, we showed that three hormonal cues, including auxin, have been recruited independently to co-ordinate branch patterning in flowering plant leafy shoots and moss gametophores (Coudert, Palubicki *et al*., 2015)^2–4^. Moreover, auxin-mediated apical dominance, which relies on local auxin production, has been proposed as a unifying molecular regulatory mechanism of branch development across land plants^5^. Whilst our previous work in the moss *Physcomitrium patens* has gathered indirect evidence supporting the notion that auxin synthesized in gametophore apices regulates branch formation at a distance^2^, direct genetic evidence for a role of auxin biosynthesis in gametophore branching control is still lacking. Here, we show that gametophore apex decapitation promotes branch emergence through massive and rapid transcriptional reprogramming of auxin-responsive genes and altering auxin biosynthesis gene activity. Specifically, we identify a subset of *P. patens TRYPTOPHAN AMINO-TRANSFERASE* (*TAR*) and *YUCCA FLAVIN MONOOXYGENASE-LIKE* (*YUC*) auxin biosynthesis genes expressed in apical and basal regions of the gametophore, and show that they are essential for branch initiation and outgrowth control. Our results demonstrate that local auxin biosynthesis coordinates branch patterning in moss and thus constitutes a shared and ancient feature of shoot architecture control in land plants.

## Introduction

Understanding the mechanisms that determine the shape of living organisms is a central goal of biology. In plants, branching patterns are a major determinant of shape diversity^6,7^. Indeed, most plant shoots have the capacity to develop indeterminately, which allows for the continuous production of new growth axes, or branches, throughout their lifetime^7^. Apical dominance, the inhibitory effect exerted by shoot apices on the initiation or outgrowth of distant lateral buds, is a fundamental regulatory mechanism of branching form development. It was first described in vascular plants and is mediated, at least partly, by the production of the phytohormone auxin at the dominant shoot apex^3,8^. Studies in mosses have shown that excision of the gametophore (*i.e*. the gametophytic leafy shoot) tip – referred to as decapitation – triggers *de novo* branch formation^2,9^, indicating that apical dominance is conserved beyond vascular plants. Using the moss *Physcomitrium patens* as a model, we have shown that decapitation-induced branching can be alleviated by the application of auxin-impregnated lanolin on the stump of decapitated gametophores^2^. We have further demonstrated the inhibitory effect of auxin on branch formation by quantifying the branching patterns of *P. patens short internodes 2-1* (*shi2-1*) mutants that lack a putative transcriptional regulator of *YUCCA FLAVIN MONOOXYGENASE-LIKE* (*YUC*) auxin biosynthesis genes^10,11^, *SHI1* over-expressors and wild-type gametophores treated with natural and synthetic auxins^2^. We have also used a computational modelling approach to investigate whether a regulatory mechanism of branch initiation by auxin-mediated apical dominance could account for observed branch distribution patterns in wild-type and mutant gametophores^2^. Whilst our model supports the notion that local auxin production in the main gametophore apex and branch apices patterns branch formation at the whole-shoot level, direct genetic evidence for local auxin production in branch apices is lacking and a role for auxin biosynthesis genes in the regulation of gametophore branching has not been demonstrated yet. More generally, the relative contribution of auxin to gametophore branching is poorly understood and non-hormonal regulatory mechanisms are yet to be identified. In this study, we took advantage of the possibility to trigger branch formation by gametophore decapitation in the moss *P. patens* to address these fundamental biological questions.

## Results

### Massive and rapid reprogramming of early auxin-responsive genes follows gametophore decapitation

To first assess the dynamics of branch emergence following decapitation, we isolated gametophores from 4-week-old wild-type colonies, cut off their apices and followed the development of branches over 48 hours. The first emerged branches were clearly visible 24 hours after decapitation (h.a.d.) and branches with well-developed leaves could be observed 48 h.a.d., suggesting that the earliest molecular events leading to branching occur within a day (Figure 1A). To capture a comprehensive overview of the transcriptional changes associated with early branch development, we next analyzed the transcriptome of whole gametophores collected 2, 6, 12 and 24 h.a.d., and used gametophores collected immediately after decapitation (0 h.a.d.) as a reference. The proportion of genes whose transcription was affected increased up to 12 h.a.d., leading to a maximum of 5028 and 4734 genes down- and up-regulated, respectively, suggesting that major transcriptional changes occur before the first branches become visible (Table S1, Figure S1 and S2). Gene ontology (GO) term analyses identified distinct enriched functional categories in both datasets (Table S2), but auxin-related GO terms were not significantly enriched, which was unexpected as published data suggest an important role for auxin in gametophore branching control^2^. To specifically address the contribution of auxin in decapitation-induced branching, we sought to directly compare the effect of auxin and decapitation at the transcriptional level. To this end, we sequenced the transcriptome of 4-week-old wild-type gametophores treated with indole-3-acetic-acid (IAA) for a short time period (30 min) and used mock-treated plants as a reference to identify early auxin-responsive genes. We found that 3652 and 4524 genes were down- and up-regulated by IAA, respectively (Table S1). By cross-referencing decapitation and IAA-regulated gene sets, we observed a striking anti-correlation between both treatments (Figure 1B). Sixty-four percent of genes down-regulated 2 h.a.d. were IAA-induced, whilst only 5% were IAA-repressed. A similar trend was observed for “6 h.a.d.” (IAA-induced genes, 61%; IAA-repressed genes, 6%) and “12 h.a.d.” (IAA-induced genes, 51%; IAA-repressed genes, 8%) gene sets. For example, both *ARF* and *AUX/IAA* auxin signalling genes, previously identified as early auxin-responsive genes^12^, responded oppositely in that all the genes but one were induced by IAA and repressed after decapitation (Figure 1C). Inversely, 46% of genes up-regulated 2 h.a.d. were IAA-repressed, whilst only 14% were IAA-induced, with a similar trend for “6 h.a.d.” (IAA-repressed genes, 49%; IAA-induced genes, 8%) and “12 h.a.d.” (IAA-repressed genes, 34%; IAA-induced genes, 11%) gene sets (Figure 1B). Thus, the opposite effect of decapitation and auxin application on the overall gametophore transcriptional response support the notion that the main gametophore apex is a major auxin source. Taken together, our data indicate that early auxin-responsive genes play a major role in the decapitation response.

**Figure 1.**
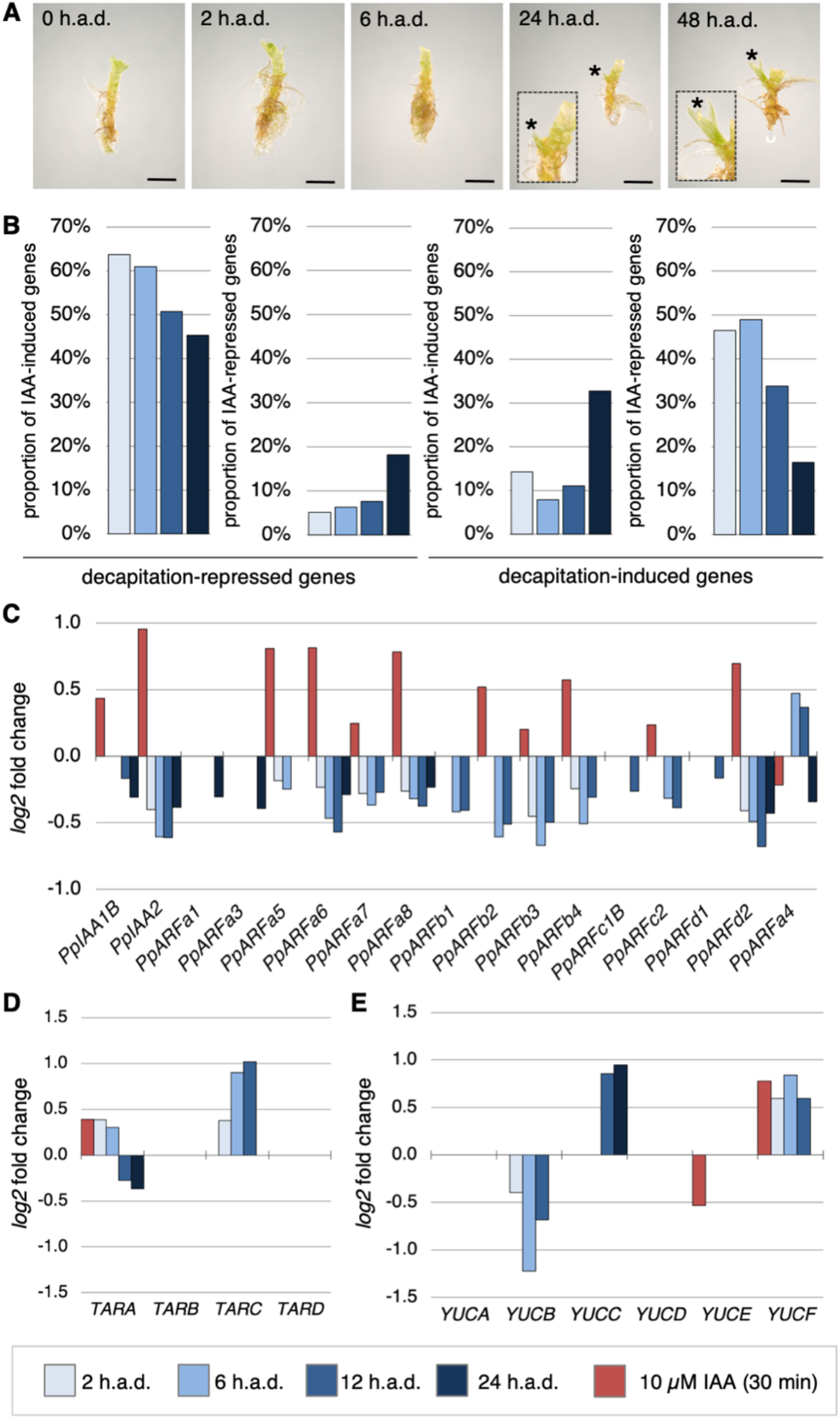
Decapitation affects the expression of early auxin-responsive genes, and auxin signalling and biosynthesis genes. (**A**) New branches (indicated with asterisks) emerged within 24 hours after gametophore decapitation. Leaves were removed before imaging to observe gametophore branches. Insets show magnifications of emerging branches. Scale bar = 1 mm. (**B**) Cross-analysis of transcriptomes of decapitated and auxin-treated gametophores showed that about two thirds of the genes repressed 2 h.a.d. were induced by a 30-minute treatment with 10 μM indole-3-acetic acid (auxin), and conversely nearly half of the genes induced 2 h.a.d. were repressed by auxin. (**C**) Bar plots showed that all but one *ARF* and *AUX/IAA* auxin signalling genes identified were repressed after decapitation and induced by auxin. (**D-E**) Expression of two *TAR* and three *YUC* auxin biosynthesis genes was affected by decapitation. The colour code shown at the bottom of the figure is used in panels B-E.

### Decapitation rapidly alters *TAR* and *YUC* auxin biosynthesis gene expression

We reasoned that our dataset would provide a reference to identify genes controlling apical dominance, both in an auxin-dependent and independent manner (see Discussion). In this context, we sought to test further the hypothesis that gametophore apices are sites of auxin biosynthesis. Auxin is derived from tryptophan (TRP)-independent and dependent biosynthetic pathways^13^. In the latter, the dominant pathway in plants, TRP is converted to indole-3-pyruvic acid (IPyA) by TRYPTOPHAN AMINO-TRANSFERASE (TAA/TAR) enzymes, that is then converted to auxin by YUCCA FLAVIN MONOOXYGENASE-LIKE (YUC) enzymes^14–17^. Auxin metabolite profiling in *PpTAR* and *PpYUC* overexpression lines, and *pptar* mutants have suggested that both steps of auxin biosynthesis are essential and identified TRP-to-IPyA conversion as a rate-limiting step in *P. patens*^13,18,19^. To determine the changes in auxin biosynthesis gene activity during decapitation-induced branching, we therefore retrieved *PpTAR* (named *TAR* hereafter for the sake of readability) and *PpYUC* (named *YUC* hereafter) expression profiles from our RNA-seq data. We identified two *TAR* and three *YUC* genes significantly differentially expressed after decapitation (Figure 1D and 1E). Specifically, *TARA* (*Pp3c21_15370V3.1*), *TARC* (*Pp3c17_6500V3.1*) and *YUCF* (*Pp3c3_20490V3.1*) were induced as early as 2 h.a.d., *YUCC* (*Pp3c1_11500V3.1*) was induced from 12 h.a.d., and *YUCB* (*Pp3c11_11790V3.1*) was repressed. We also found that the expression of *TARA, YUCE* (*Pp3c13_21970V3.1*) and *YUCF*, but not *TARC, YUCB* and *YUCC*, was affected by exogenous auxin, suggesting that auxin may to some extent regulate its own biosynthesis.

### *TAR* and *YUC* genes are active in the tip, branch apices and basal portion of gametophores

To determine the expression domain of *TARA, TARC, YUCB, YUCC* and *YUCF* genes, we analysed the activity of transcriptional fusions between corresponding promoter regions and a GFP-GUS chimeric protein transformed in a wild-type background (Figure S3). Two to three independent transgenic lines were observed for each genetic construct (Table S4). We found that the spatial expression domains of *TAR* and *YUC* promoters largely overlapped in whole gametophores. Beta-glucuronidase (GUS) activity was detected at the main apex, at the base, notably in rhizoid cells, and in discrete regions of the stem corresponding to initiating branch apices (Figure 2A, 2D, 2G, 2J and 2M). At the main apex, *TAR* promoter activity was observed in the apical cell and emerging leaves, as reported previously^18^, and both *TAR* and *YUC* promoters were highly active in axillary hair cells surrounding the gametophore apical cell, reminiscent of the expression pattern of other auxin-related genes like *PpSHI1* and *PpSHI2*^10^ (Figure 2P and 2Q). Promoter activity of *YUCF,* and to a lesser extent *YUCB* and *YUCC*, was also detected in axillary hair cells elsewhere on the stem (Figure 2M, 2R). This indicates that the expression of *TARA* and *TARC* is more restricted in space than that of *YUCB, YUCC* and *YUCF*, and it is therefore unlikely that auxin biosynthesis occurs in leaf axils in the absence of *TARA* or *TARC* activity.

**Figure 2.**
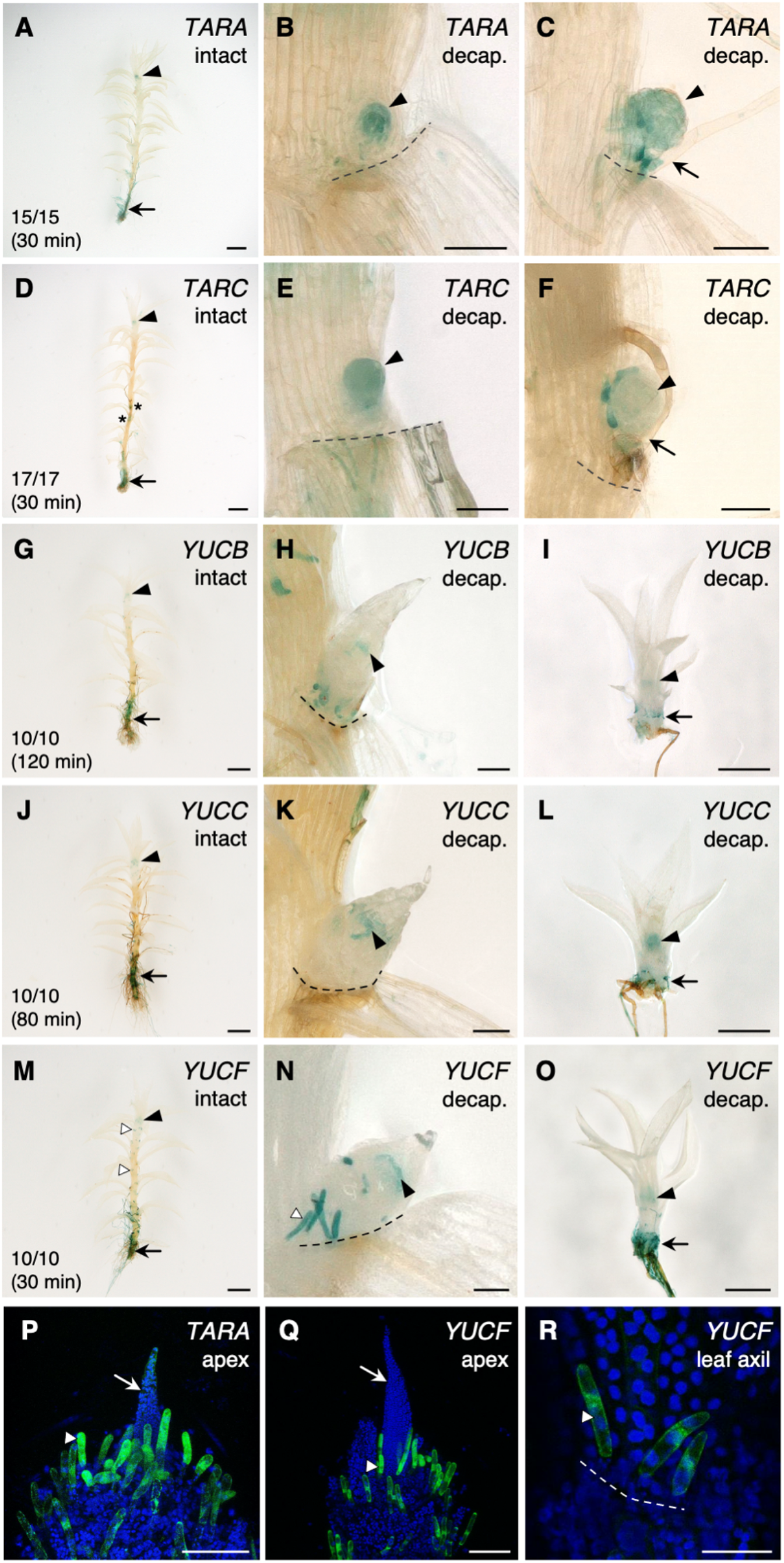
*TAR* and *YUC* genes are expressed in meristematic and basal regions of *Physcomitrium* gametophores. (**A-O**) GUS staining of *TARA::GFP-GUS* (A-C), *TARC::GFP-GUS* (D-F), *YUCB::GFP-GUS* (G-I), *YUCC::GFP-GUS* (J-L) and *YUCF::GFP-GUS* (M-O) transgenic lines revealed that *TAR* and *YUC* genes are active in largely overlapping expression domains. Gametophore were either observed before (“intact”) or after (“decap.”) apex excision. In intact gametophores, *TARA* and *TARC* are expressed in the main apex (black arrowheads), in emerging branches (asterisks) and in the basal region (black arrows) (A, D). Following decapitation, *TARA* and *TARC* expression is detected in initiating lateral branches (black arrowheads) but not in rhizoids (black arrows) (B, C, E, F). In intact gametophores, *YUCB, YUCC* and *YUCF* are expressed in the main apex (black arrowheads) and in the basal region (black arrows). *YUCF* expression is also observed in axillary hairs in leaf axils (white arrowheads) (G, J, M). Following decapitation, *YUCB, YUCC* and *YUCF* expression is first detected at a later stage than *TARA* and *TARC*. GUS staining was found in axillary hairs at the apex of newly formed branches (black arrowheads) and, at a later stage, in branch rhizoids (black arrows) (H, I, K, L, N, O). (**P-R**) GFP signal in *TARA::GFP-GUS* and *YUCF::GFP-GUS* lines was strongest in hairs (white arrowheads) located at the gametophore apex (P-Q), and/or in leaf axils (R). Emerging leaves are indicated with white arrows. Dashed lines mark the boundary between the stem and the detached leaf (B, C, E, F, H, K, N, R). Gametophores were soaked in GUS staining solution for times specified in panels A, D, G, J and M. The scale bars represent 1 mm in A, D, G, J and M, 500 μm in I, L, and O, 100 μm in B, C, E, F, H, K, P and Q, and 50 μm in N and R.

To determine how decapitation affected the activity of selected *TAR* and *YUC* transcriptional reporters, gametophores were collected 24 hours after apex excision and stained for GUS activity, thereby leaving enough time for new branches to initiate and grow out. The intensity of GUS accumulation patterns remained overall similar to intact controls, except for the *YUCB* promoter that seemed more weakly expressed after decapitation, likely reflecting its transcriptional repression (Figure 1E). The spatial distribution of these patterns was changed in leaf axils where we observed an activation of *TARA, TARC, YUCB, YUCC* and *YUCF* expression associated with the formation of new branches. The activity of *TARA* and *TARC* promoters was detected from the earliest stage of branch formation, throughout the branch initium (Figures 2B, 2C, 2E and 2F). In contrast, *YUCB, YUCC* and *YUCF* promoter activity were first detected at a later stage when the first leaves were clearly visible, and mostly in hair cells of the branch apical region (Figures 2H, 2I, 2K). At a more mature stage of branch development, *i.e*. when leaves and brown rhizoids were well-developed, *TARA, TARC, YUCB, YUCC* and *YUC*F genes were active in overlapping spatial domains mirroring those described previously in whole gametophores (Figures 2I, 2L and 2O, data not shown for *TARA* and *TARC*). The temporal delay between *TAR* and *YUC* gene activation and the differences in their spatial expression domains at early stages of branch initiation suggest that IPyA might be transported from the meristematic cells where it is synthesized to adjacent hair cells to be converted to auxin by YUC enzymes. Alternatively, these differences might be explained by the short-range movement of *TAR* or *YUC* mRNA, or corresponding proteins; the involvement of *YUC* genes co-expressed with *TARA* and *TARC* but not differentially expressed in our RNA-seq dataset, such as *YUCE* (Figure S4); or unmask a putative YUC-independent function of IPyA. Together, the above data suggest that TRP-dependent auxin biosynthesis occurs primarily in the main apex and the basal portion of gametophores, and resumes locally in leaf axils when new branches initiate.

### *TAR* genes prevent systematic branch initiation

To investigate the function of *TARA* and *TARC* genes in gametophore branching, we grew *tara, tarc* and *tarac* knock-out mutants^18^ for 5 or 7 weeks and quantified their branching patterns, as done previously^2^. We found that *tara* single mutants had similar phenotypes to wild-type control plants. There was moderate evidence^20^ that *tarc* branch density was lower than control plants in older gametophores, indicating that *tarc* single mutants were also very similar to wild-type (Figures 3A-3C, 3E-3G and 3K-3N). In contrast, there was strong evidence that *tarac* double mutants had a shorter apical inhibition zone and inter-branch distance, and an increased branch density with respect to wild-type plants, suggesting a reduced apical dominance and lateral inhibition from branch meristems (Figure 3D, 3H, 3K, 3L and 3O). We also observed that the position of the lowermost branch was closer to the gametophore base in *tarac* mutants than in wild-type (Figure 3P), and the number of branches in the three most basal metamers was much higher in *tarac* mutants (n = 23 branches) than in *tara* (n = 2), *tarc* (n = 2) or wild-type (n = 5) gametophores, suggesting that local *TAR* activity prevents branching at the gametophore base (Figures 3A-3D). Despite the overall increase in branch number in *tarac* mutants, we observed strong disparities in the branch distribution pattern between gametophores. Whilst some gametophores bore branches on up to six consecutive metamers (see gametophore #1 in Figure 3D), which has never been reported elsewhere before, others had few or no visible branches at all, which was rather counter-intuitive (Figure 3D). However, on closer inspection of *tarac* gametophores, we noticed that tiny buds were nested in nearly every single leaf axil lacking a visible branch (Figure 3I-3J). This suggests that branch initiation was almost systematically de-repressed in *tarac* mutants while subsequent outgrowth was inhibited. Consistent with observed *TARA* and *TARC* expression patterns in whole gametophores and emerging branches, our data indicate that both genes act redundantly in the apices and basal portion of gametophores and their activity is necessary to prevent branch initiation and co-ordinate branch patterning.

**Figure 3.**
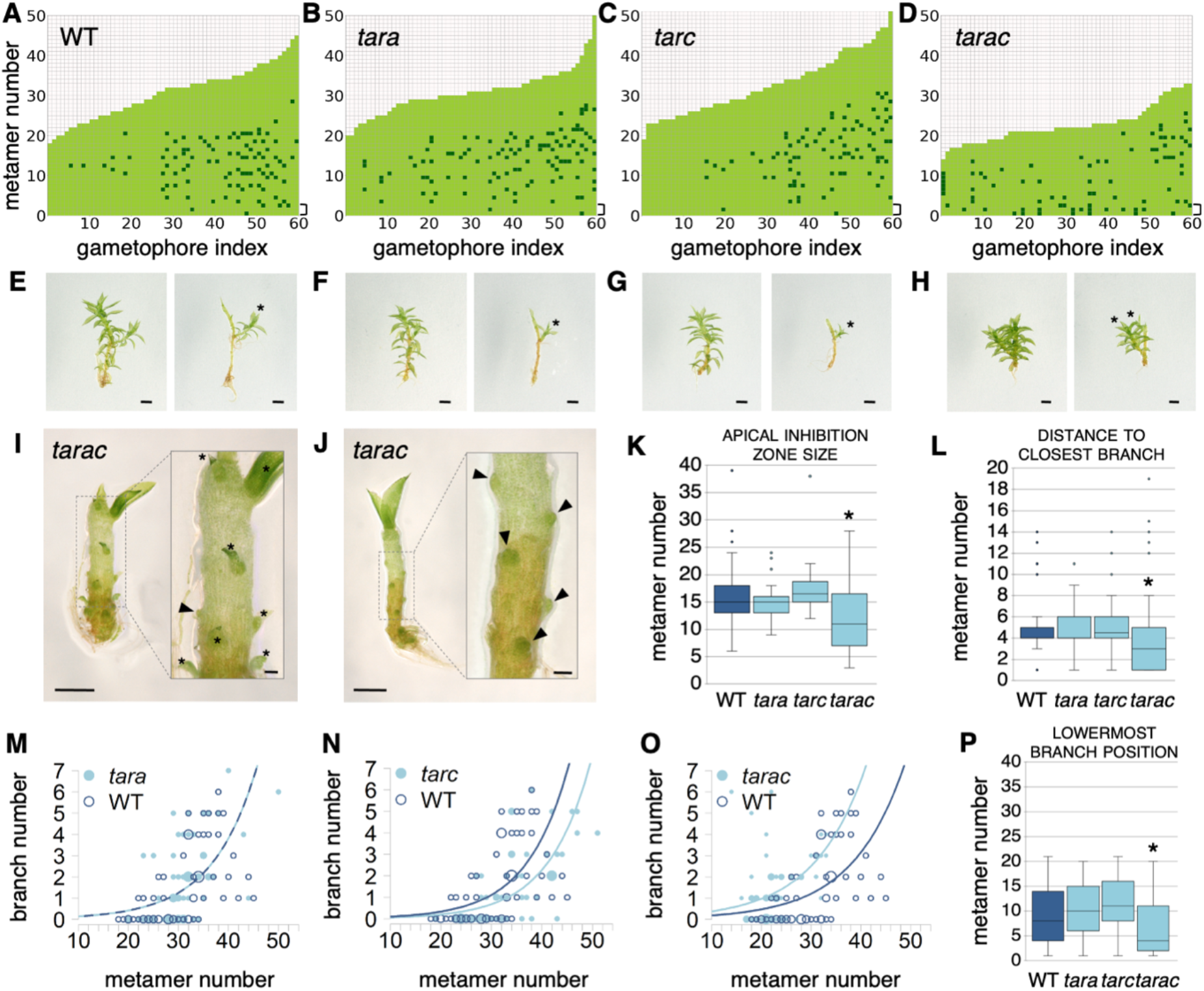
*TARA* and *TARC* genes prevent branch initiation and pattern branch distribution along the gametophore axis. (**A-D**) Branching patterns in WT, *tara*, *tarc* and *tarac* mutants are presented as in Coudert, Palubicki *et al*. (2015)^2^. Brackets indicate the three lowermost metamers. (**E-H**) WT (E), *tara* (F), *tarc* (G) and *tarac* (H) gametophores before (left) and after (right) removing leaves, with asterisks marking branches. Scale bars represent 1 mm. (**I-J**) *tarac* gametophores with leaves removed revealed lateral buds with no visible leaves (arrowheads) and tiny branches with few visible leaves (asterisks), which is never observed in WT in which branches initiate *de novo* and grow out with no dormancy period. Scale bars represent 500 μm in the main panels and 100 μm in the insets. (**K**) There was strong evidence^20^ that the apical inhibition zone was reduced in *tarac* double mutants, but not in *tara* or *tarc* single mutants, compared with wild-type control (Wilcoxon rank sum test with continuity correction different from WT, *p-value ≤ 0.05; WT versus *tara*, p-value = 0.35; WT versus *tarc,* p-value = 0.07; WT versus *tarac,* p-value = 0.005). (**L**) There was very strong evidence that the distance to the closest branch, a measurement of branch spacing, was significantly reduced in *tarac* double mutants, but not in *tara* or *tarc* single mutants, compared with wild-type control (Wilcoxon rank sum test with continuity correction different from WT, *p-value ≤ 0.05; WT versus *tara*, p-value = 0.92; WT versus *tarc,* p-value = 0.14; WT versus *tarac,* p-value = 0.0009). (**M-O**) Bubble plots showed that branch number at a given length is increased in *tarac*, slightly decreased in older *tarc* gametophores, and unchanged in *tara*, compared with WT. Gametophore length is represented as the number of metamers and the bubble area is proportional to the number of gametophores with the same branch number (*B*) at a given length (*L*). The data were over-dispersed, and so negative binomial regression was used to test whether and how the relationship between branch number and gametophore length differed between mutants and WT (see ‘Material and methods’). For (M) the best-fitting relationship indicated no difference between *tara* and WT (p-value = 0.39; 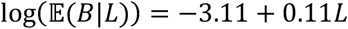 for both treatments). For (N) the best-fitting relationship indicated WT was larger than *tarc* at all lengths (p-value = 0.01; 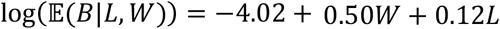, where *W* = 1 for WT). For (O) the best-fitting relationship indicated WT was smaller than *tarac* at all lengths (p-value=0.006; 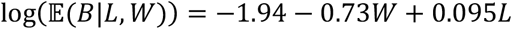, where *W* = 1 for WT). (**P**) The position of the lowermost branch was closer to the gametophore base in *tarac* than in *tara* or *tarc* in comparison with WT (Wilcoxon rank sum test with continuity correction different from WT, *p-value ≤ 0.05; WT versus *tara*, p-value = 0.30; WT versus *tarc*, p-value = 0.12; WT versus *tarac*, p-value = 0.007).

### *YUC* genes suppress branching

Then, to determine the role of decapitation-induced *YUC* genes in gametophore branching, we grew *yucc, yucf* and *yuccf* knock-out mutants^18^ for 5 or 7 weeks and quantified their branching patterns^2^ (Figure S5). Whilst the apical inhibition zone size was increased in *yucc* gametophores, the distance to the closest branch, the position of the lowermost branch of *yucc* and *yucf* gametophores were not changed with respect to wild-type controls (Figure 4A-4C, 4E-4G, 4I-4M). We observed slight perturbations of the branch distribution patterns in *yucc* and *yucf* single mutants, mostly explained by the branching patterns of older gametophores (Fig 4L-M). In contrast, we found strong evidence that *yuccf* double mutants had a reduced apical inhibition zone and inter-branch distance, and moderate evidence of an increased branch density with respect to wild-type plants (Figure 4D, 4H, 4I-4K, 4N). Moreover, the position of the lowermost branch was closer to the gametophore base in *yuccf* double mutants than in wild-type (Figure 3P), and the number of branches in the three most basal metamers was higher in *yuccf* mutants (n = 15 branches) than in *yucc* (n = 11), *yucf* (n = 6) or wild-type (n = 3) gametophores. Unlike *tarac* mutants, we did not observe emerging buds in the leaf axils of *yuccf* gametophores. However, we sometimes noticed supernumerary branches growing from the base of lateral branches in *yucc* mutants, consistent with *YUCC* expression at the base of branches (Figure 4F) and indicating a local inhibitory effect of *YUCC*. Moreover, branches seemed less developed in both single and double *yuc* mutants than in wild-type plants. Together, the above data coincide with observed expression patterns of *YUC* transcriptional reporters and indicate that *YUCC* and *YUCF* act redundantly to inhibit branch formation but promote subsequent growth, hence supporting the hypothesis of a local role of auxin biosynthesis in branch patterning.

**Figure 4.**
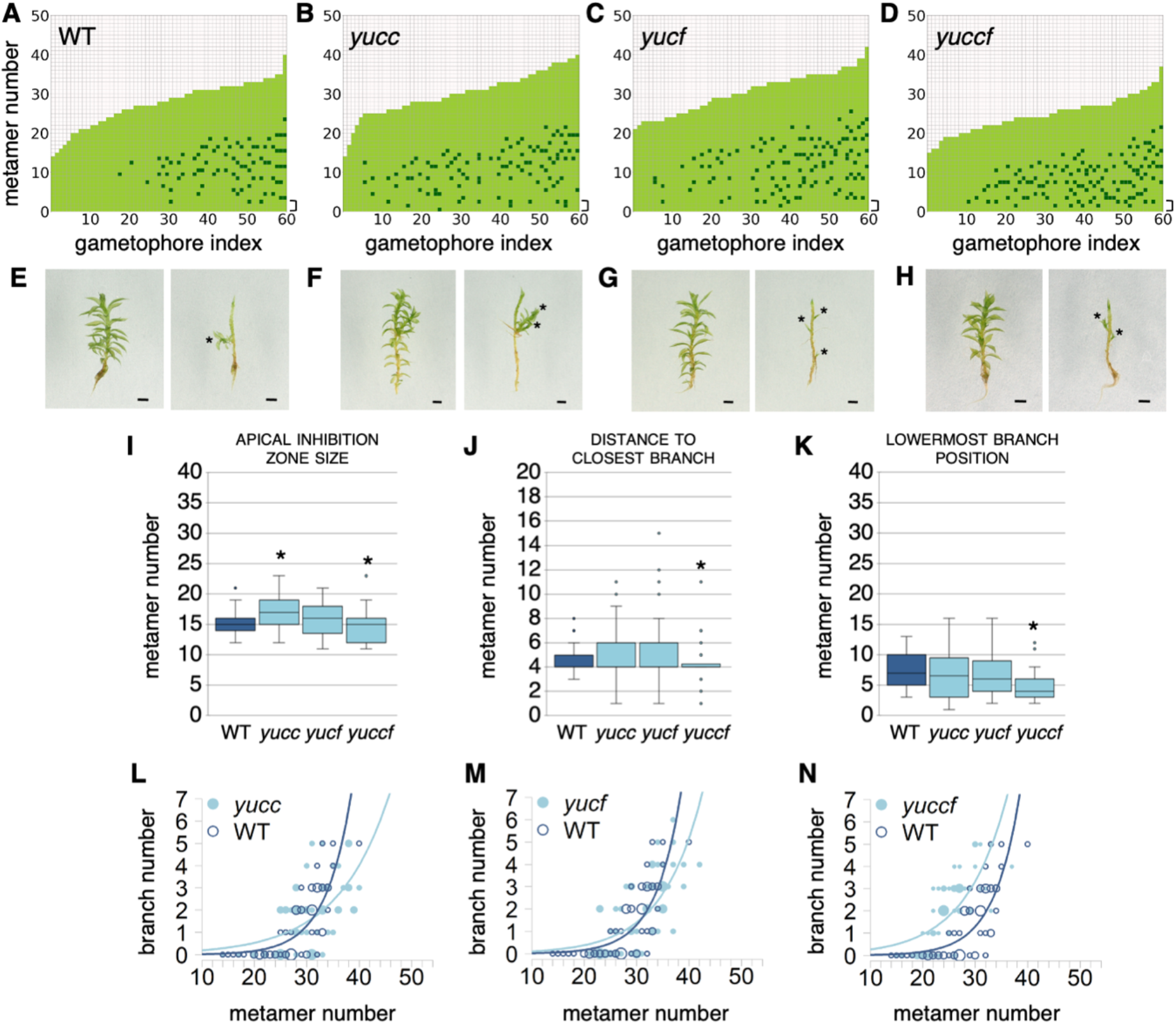
*YUCC* and *YUCF* genes repress branching. (**A-D**) Branching patterns in WT, *yucc*, *yucf* and *yuccf* mutants. Brackets indicate the three lowermost metamers. (**E-H**) WT (E), *yucc* (F), *yucf* (G) and *yuccf* (H) gametophores before (left) and after (right) removing leaves, with asterisks marking branches. Scale bars represent 1 mm. (**I**) There was strong and moderate evidence that the apical inhibition zone was different in *yucc* and *yuccf* mutants, respectively, compared with wild-type control (Wilcoxon rank sum test with continuity correction different from WT, *p-value ≤ 0.05; WT versus *yucc*, p-value = 0.003; WT versus *yucf*, p-value = 0.82; WT versus *tarac*, p-value = 0.05). (**J**) There was strong evidence that the distance to the closest branch, a measurement of branch spacing, was reduced in *yuccf* double mutants, but not in *yucc* or *yucf* single mutants, compared with wild-type control (Wilcoxon rank sum test with continuity correction different from WT, *p-value ≤ 0.05; WT versus *yuc*, p-value = 0.77; WT versus *yucf*, p-value = 0.76; WT versus *yuccf*, p-value = 0.01). (**K**) There was very strong evidence that the position of the lowermost branch was closer to the gametophore base in *yuccf* than in *yucc* or *yucf* in comparison with WT (Wilcoxon rank sum test with continuity correction different from WT, *p-value ≤ 0.05; WT versus *yucc*, p-value = 0.49; WT versus *yucf*, p-value = 0.34; WT versus *yuccf*, p-value = 0.0006). (**L-M**) Bubble plots showed that branch number responded differently to length in all three mutants compared with WT. Gametophore length is represented as the number of metamers and the bubble area is proportional to the number of gametophores with the same branch number *(B)* at a given length (L). The data were not over-dispersed, and so Poisson regression was used to test whether and how the relationship between branch number and gametophore length depended on treatment (see ‘Material and methods’). The best-fitting relationship included an interaction for all three mutants, indicating the nature of the difference in the number of branches between WT and mutant depended upon the length (in (L), p = 0.002, 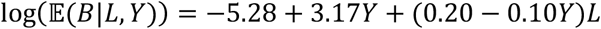, where *Y* = 1 for *yucc*; in (M) p = 0.03, 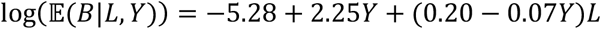, where *Y* = 1 for *yucf*; in (N) p = 0.02, 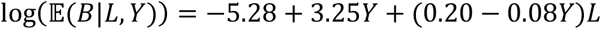, where *Y* = 1 for *yuccf*).

## Discussion

This work primarily aimed to determine the relative importance of auxin in gametophore branching and test the hypothesis that apically synthesized auxin patterns branch distribution. Using an RNA-seq approach, we show that early auxin-responsive genes have a major contribution to the transcriptional response of gametophores to decapitation. We also provide evidence that several *TAR* and *YUC* genes, together responsible for the enzymatic conversion of tryptophan to auxin, are active in the main apex and branch apices, and the base of gametophores^13,18^, and identify a prominent and dual role for *TARA* and *TARC* in branch initiation and growth. First, our data show that apical auxin sources suppress branch initiation at a distance. Indeed, observed perturbations of branching patterns in *tarac* and *yuccf* mutants (*i.e*. the reduction of the apical inhibition zone size and distance between closest branches, and the increase of branch density) qualitatively match the predicted outcome of our published computational model of gametophore branching control by hormonal cues where a reduction of auxin production in the main and lateral apices promotes branching^2^. Interestingly, the model has also identified that a basal inhibitory cue is locally required to prevent branching, and genetic experiments previously led us to propose that this cue is a *CCD8*-derived molecule belonging to the strigolactone family of compounds^2^. Expression patterns of selected *TAR* and *YUC* transcriptional reporters indicate that auxin biosynthesis occurs in this region, and basal branching is strongly increased in *tarac* and *yuccf* double mutant gametophores, suggesting that auxin may act concomitantly with the *CCD8*-derived cue to fulfil this inhibitory role. Determining whether and how these two hormonal cues interact will require further investigation. Second, our data indicate that an adequate level of *TAR* and *YUC* activity at branch initiation sites is essential to drive branch outgrowth. Branches were noticeably shorter in *tarac* mutants, and likely also in *yuccf* mutants, than in wild-type. These observations corroborate the finding that *tarac* gametophores have stunted growth^18^ and suggest that low auxin biosynthesis in gametophore apices hinders apical cell proliferative activity. It has previously been reported that excessive auxin accumulation in gametophore tissues may also disrupt apical cell function and suppress phyllid (*i.e*. the gametophytic leaf) emergence at the apex^21^, which supports the idea that cellular auxin homeostasis must be tightly controlled to ensure proper gametophore growth. Third, despite two whole-genome duplications in *P. patens* and their possible consequences on gene redundancy^22^, it must be stressed that deleting only 2 out of 4 *TARs*, and 2 out of 6 *YUCs* was sufficient to reveal the function of both gene families in gametophore branching control. Nevertheless, it cannot be excluded that other *TAR* or *YUC* genes may also be involved in this process.

From an evolutionary perspective, TAR-YUC-mediated auxin biosynthesis originated in streptophyte algae and is conserved in land plants^23,24^. The genome of the liverwort *Marchantia polymorpha* contains only one functional *TAA* gene, named *MpTAA,* and two YUC genes, named *MpYUC1* and *MpYUC2*^25^. Similar to our observations in moss gametophores, *MpYUC2* expression domain is broader than that of *MpTAA* in the liverwort thallus (*i.e*. the gametophytic body), and both domains overlap in meristematic notches that contain the apical stem cells at the origin of thallus tissues. Although thallus branching occurs by notch dichotomy, a type of apical branching that is developmentally distinct from gametophore branching^7,26^, *Marchantia taa* and *yuc* knock-down plants also display hyperbranching phenotypes. Beyond bryophytes, various studies in the flowering plant *Arabidopsis thaliana* have shown that *AtTAA1/WEI8, AtTAR2, AtYUC1*, *AtYUC4* and *AtYUC6* genes are expressed in discrete and partially overlapping domains of the shoot apical meristem^14,15^. Double and triple mutants in these genes, including *yuc1yuc4, yuc1yuc4yuc6* and *taa1tar2* allele combinations, show various degrees of decreased apical dominance reflected by an enhanced production of branches. Together, this suggests that lateral and apical branching modes evolving separately in flowering plant sporophytes, and moss and liverwort gametophytes, are underpinned by a shared and ancient molecular mechanism driving local auxin biosynthesis in shoot meristematic structures, which represents a novel discovery of deep homology in land plants^21,27,28^.

The functional analysis of *PpTAR* and *PpYUC* auxin biosynthesis genes demonstrated that our RNA-seq approach has the potential to identify novel regulators of gametophore branching. Moreover, the relatively small differences in gene expression detected at the whole gametophore level (see for example Figure 1D-E) likely under-estimate the magnitude of regulatory changes occurring locally, since genes underpinning branch initiation are expressed in restricted spatial domains. In the future, our dataset will help us to decipher how auxin is integrated at the molecular level to regulate branching. For instance, GO analyses revealed a significant enrichment of the “DNA binding transcription factor activity” term in gene sets up- and down-regulated after decapitation (Table S2). Members of the *APETALA2/ETHYLENE RESPONSIVE FACTOR* (*AP2/ERF*) family represented about two thirds of identified DNA binding transcription factor coding genes and included putative conserved regulators of meristematic function, such as *STEMIN1* (*Pp3c1_27440v3.1*), known to promote stem cell formation through local depletion of a repressive chromatin mark, and *Pp3c26_9880v3.1*, the closest homologue to *Solanum lycopersicum LEAFLESS* and *A. thaliana PUCHI* and *DORNRÖSCHEN-LIKE*, which are regulated by auxin and involved in meristem identity and lateral organ initiation control^29–32^. We also found that GO terms related to “trehalose metabolism” were significantly enriched in the “up-regulated 2 h.a.d.” gene set. Interestingly, trehalose 6-phosphate (T6P) metabolic genes regulate branching and apical dominance in flowering plants^33–35^. For example, shoot decapitation in *Pisum sativum* (pea) triggers an increase of T6P levels in a few hours, which contributes to promoting branch outgrowth^33^. T6P is synthesized from uridine diphosphate-glucose and glucose-6-phosphate by the activity of T6P SYNTHASES (TPS) and converted to trehalose by T6P PHOSPHATASES (TPP), which is then hydrolysed into glucose by TREHALASES (TRE). Transgenic *Arabidopsis thaliana* plants with increased TPP activity in axillary buds have reduced T6P levels and delayed branch outgrowth, whilst plants with increased levels of T6P in the vasculature have enhanced branching^34^. Moreover, mutations in *Zea mays RAMOSA3* (*RA3*) and its paralog *ZmTPP4* that both encode TPP enzymes, lead to increased inflorescence branching^35,36^. In *Physcomitrium*, we found that all *PpTPS* genes were repressed after decapitation, whilst *PpTPP* and *PpTRE* genes were either repressed or induced (Figure S6), suggesting that T6P levels are dynamically regulated during decapitation-induced branching. Besides, we found that *TPS* genes were IAA-induced and decapitation-repressed, and GO terms related to trehalose metabolism were associated with genes both induced by auxin and up-regulated after decapitation (Table S3), consistent with *TPP* and *TRE* expression profiles 2 and 6 h.a.d. (Figure S6). This suggests that apically-produced auxin regulates the activity of T6P metabolic genes in the gametophore, which has not been evidenced yet in flowering plants. Further studies will be needed to determine the biological function of T6P metabolic genes and AP2/ERF DNA-binding transcription factors, and explore their crosstalk with auxin in the moss *P. patens*. Finally, it must be stressed that our dataset contains a large number of differentially expressed genes with uncharacterized function, representing an untapped reservoir of novel putative regulators of shoot architecture.

## Material and methods

### *Physcomitrium patens* plant growth and transformation

The Reute^37^ strain of *Physcomitrella patens* was used as the wild-type (WT) background for generating all the transgenic lines, the Gransden strain being largely sterile in laboratory conditions. Moss colonies were initiated from 1 mm^2^ spot cultures and cultivated in sterile Magenta GA-7 tissue culture vessels (Bioworld, Dublin, OH, USA) on BCDAT medium (250 mg/L MgSO_4_.7H_2_O, 250 mg/L KH_2_PO_4_ (pH 6.5), 1010 mg/L KNO_3_, 12.5 mg/L, FeSO_4_.7H_2_O, 0.001% Trace Element Solution (0.614 mg/L H_3_BO_3_, 0.055 mg/L AlK(SO_4_)_2_.12H_2_O, 0.055 mg/L CuSO_4_.5H_2_O, 0.028 mg/L KBr, 0.028 mg/L LiCl, 0.389 mg/L MnCl_2_.4H_2_O, 0.055 mg/L CoCl_2_.6H_2_O, 0.055 mg/L ZnSO_4_.7H_2_O, 0.028 mg/L KI and 0.028 mg/L SnCl_2_.2H_2_O), 0.92 g/L di-ammonium tartrate (C_4_H_12_N_2_O_6_) and 8 g/L agar with CaCl_2_ added to a 1 mM concentration after autoclaving, at 23°C under a 16h light/8h dark cycle, at 50-150 μmol.m^-2^.s^-1^ in MLR-352 growth cabinets (PHCbi, Etten-Leur, The Netherlands).

### Decapitation and auxin treatment

For pharmacological treatments, a 70% ethanol solution containing 100 mM indole-3-acetic acid (IAA) (Merck KGaA, Darmstadt, Germany) was diluted 10,000 times in water to reach a final concentration of 10 μM IAA. A 70% ethanol solution was diluted 10,000 times in water and used as a mock control. Moss cultures were soaked in diluted solutions for 30 minutes before tissue harvest. For decapitation, gametophores were teased apart from moss colonies with thin tip tweezers, planted in an upright position on BCDAT petri dishes and their apical portion was cut off with micro-scissors, as described in Coudert, Palubicki *et al*. (2015)^2^.

### Generation of *YUC* transcriptional reporters and *yuc* mutants

All the transgenic lines used in this study are listed in Table S4. Generation and confirmation of the transcriptional reporter lines *TARA::GFP-GUS, TARC::GFP-GUS*, and *YUCF::GFP-GUS*, as well as the knockout lines *tara*, *tarc, tarac, yucc*, and *yucf* have been previously described^18^. To produce the *YUCB::GFPGUS* reporter construct pMT244 (Figure S3), a 2090 bp *YUCB* promoter fragment was amplified from gDNA with primers SS235/SS236 and trimmed with *BamHI/NcoI*. The resulting fragment was cloned between the *BamHI/NcoI* sites of pMT211, a vector allowing promoters to be cloned ahead of a GFP-GUS reporter gene for subsequent integration into the neutral *Pp108* locus^38^. Similarly, to produce the *YUCC::GFPGUS* reporter construct pMT251 (Figure S3), a 1767 bp *YUCC* promoter fragment was amplified from gDNA with primers SS237/SS238, trimmed with *Bam*HI/*Bsp*HI, and cloned between the same two sites in pMT211. Both reporter constructs were linearized with *Sfi*I before they were transformed into WT protoplast as previously described^39^. Stable transformants were selected in the presence of 50 μg.ml^-1^ hygromycin (Duchefa H0192; Haarlem, the Netherlands). Correct integration was confirmed by PCR (Figure S3). For primer sequences, see Table S5. The *yuccf* double knockout lines were produced by a sexual cross of the confirmed single mutants *Ppyucc-2* and *Ppyucf-1* described previously^18^, according to the method presented in Thelander *et al*.^38^. Double knockout lines were confirmed by PCRs demonstrating the loss of internal gene sequences (Figure S5).

### GUS staining and plant imaging

*Physcomitrium patens* gametophores were isolated from colonies grown on BCDAT and incubated at 37°C in a 100 mM phosphate buffer (pH 7) with 10 mM Tris (pH 8), 1 mM EDTA (pH 8), 0.05% Triton X-100, 5mM potassium ferricyanide, 5mM potassium ferrocyanide and 2 mM X-GlcA (5-Bromo-4-chloro-3-indolyl-β-D-glucuronic acid), using times indicated in Figure 2. Tissues were de-stained in 70% ethanol and imaged with a Keyence VHX-900F digital microscope with a 5-50 X or a 50-200 X objective. Plants not subject to GUS staining were imaged in the same conditions.

### Definition of branches in *Physcomitrium patens*

According to Coudert *et al*.^1^, a module is defined as “a portion of gametophore arising from a single apical cell”. Here, we consider that the main gametophore axis corresponds to a Class I module, and gametophore branches correspond to Class II lateral modules.

### RNA-seq data production and analysis

For each biological replicate and condition, RNA were pooled from 5-10 gametophores (Gransden^40^ strain). Three independent biological replicates were produced. Total RNA was extracted using an RNeasy Plant Mini Kit (Qiagen, Hilden, Germany) and treated with DNAse according to the supplier’s instructions. RNA-seq libraries were made using the TruSeq_Stranded_mRNA_SamplePrep_Guide_15031047_D protocol (Illumina, California, U.S.A.). The RNA-seq samples have been sequenced in paired-end (PE) with a sizing of 260 bp and a read length of 75 bases, using an Illumina NexSeq500 technology (IPS2, POPS platform, Gif-sur-Yvette). 24 samples were processed by lane of NextSeq500 using individual bar-coded adapters, generating approximately 30 millions of PE reads per sample. All experimental steps, from growth conditions to bioinformatic analyses, were deposited in the CATdb database^41^ (http://tools.ips2.u-psud.fr/CATdb/, ProjectID NGS2017_09_Moss1) according to the MINSEQE ‘minimum information about a high-throughput sequencing experiment’ (https://doi.org/10.25504/FAIRsharing.a55z32). To facilitate comparisons, all samples were processed similarly from trimming to count. RNA-Seq pre-processing included trimming library adapters and performing quality controls. Raw data (fastq) were trimmed with fastx toolkit for Phred Quality Score Qscore >20, read length >30 bases, and ribosome sequences were removed with sortMeRNA^42^. The mapper Bowtie^43^ (version 2) was used to align reads against the *Physcomitrium* (*Physcomitrella*) *patens* transcriptome (with local option and other default parameters). The 32926 genes were extracted from Phytozome database (*Physcomitrella patens* transcripts v3.1 gene model). The abundance of each gene was calculated by a local script which parses SAM files and counts only paired-end reads for which both reads map unambiguously one gene, removing multi-hits. According to these rules, around 82,7% of PE reads were associated to a gene, 5-6% PE reads were unmapped and 12-13% of PE reads with multi-hits were removed. Choices for the differential analysis were made based on the article by Rigaill *et al*^44^. Genes which did not have at least one read after a counts per million (CPM) normalization in at least one half of the samples were discarded. Library size was normalized using the TMM method and count distribution was modeled with a negative binomial generalized linear model. Dispersion was estimated by the edgeR method^45^ (version 1.12.0) in the statistical software ‘R’ (http://www.R-project.org, version 2.15.0). Expression differences were compared between “0 h.a.d.” and “2 h.a.d.”, “6 h.a.d.”, “12 h.a.d.” or “24 h.a.d.” conditions for the “decapitation experiment”, and between samples “mock” and “10μM IAA” conditions for the “auxin experiment”, using likelihood ratio test and p-values were adjusted by the Benjamini-Hochberg procedure to control FDR. A gene was considered as differentially expressed if its adjusted p-value was lower than 0.05. Venn diagrams were generated using InteractiVenn^46^. Gene Ontology (GO) term enrichment analyses were performed with ShinyGO (version 0.66)^47^.

### RNA-seq data availability

RNA-seq datasets are available in the international repository GEO (Gene Expression Omnibus^48^, http://www.ncbi.nlm.nih.gov/geo) under the project identifier GSE188843 (token = ctslsswyfjchdcp).

### Sample-size estimation and replicates

For data shown in Figure 1, three independent biological replicates were produced for each condition, and each replicate corresponded to RNA extracted from 5 to 10 gametophores. For Figure 2, two or three independent transgenic lines were analysed for *TAR* and *YUC* transcriptional reporters. Independent lines transformed with the same genetic construct showed similar GUS staining patterns. Numbers in panels A, D, G, J and M indicate the proportion of gametophores with a GUS staining pattern similar to the picture. For Figure 3, data shown in panels A-D were obtained from 30 gametophores collected 5 weeks after protonema inoculation and 30 gametophores collected 7 weeks after protonema inoculation, metrics in panels K, L and P could be calculated only for gametophores with more than one branch, data in panels M-O correspond to all gametophores. For Figure 4, data shown in panels A-D were obtained from 30 gametophores collected 5 weeks after protonema inoculation and 30 gametophores collected 7 weeks after protonema inoculation, data in panels I-K could be calculated only for gametophores with more than one branch, data in panels L-N correspond to all gametophores.

### Statistical analyses

Generalised linear modelling was used to test whether the relationship between branch number and gametophore length depended on genotype (File S1). Poisson or negative binomial regression was used, depending on whether the data were over-dispersed, *i.e*. whether the data were more variable than would be expected under a Poisson model. The most complex fitted models were of the form 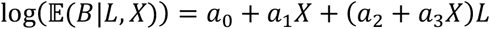, in which *a*_0_, *a*_1_, *a*_2_ and *a*_3_ are coefficients, *L* is the length, *X* is an indicator variable depending on genotype (*i.e. X* took values of either 0 or 1 depending upon whether the mutant or WT was considered), and 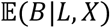 is the expected branch number (*B*) at any given pair of values of *L* and *X*. Analysis of deviance and backwards stepwise elimination was used to find the minimal model that received support from the experimental data. A mutant was considered different from WT if either the interaction term (*a*_3_) or the term corresponding to genotype (*a*_1_) remained in the minimal model, *i.e*. if the expected branch number was affected by the value of *X*. The choice of whether to use a Poisson and negative binomial model was made via a formal test of over-dispersion for the Poisson version of the model with an interaction, using the function overdispersion() in the package AER^49^. The terms “weak”, “moderate”, “strong” and “very strong evidence” reflected the translation of p-values into the language of evidence proposed by Muff *et al*.^20^

## Supplementary data

**Figure S1.**
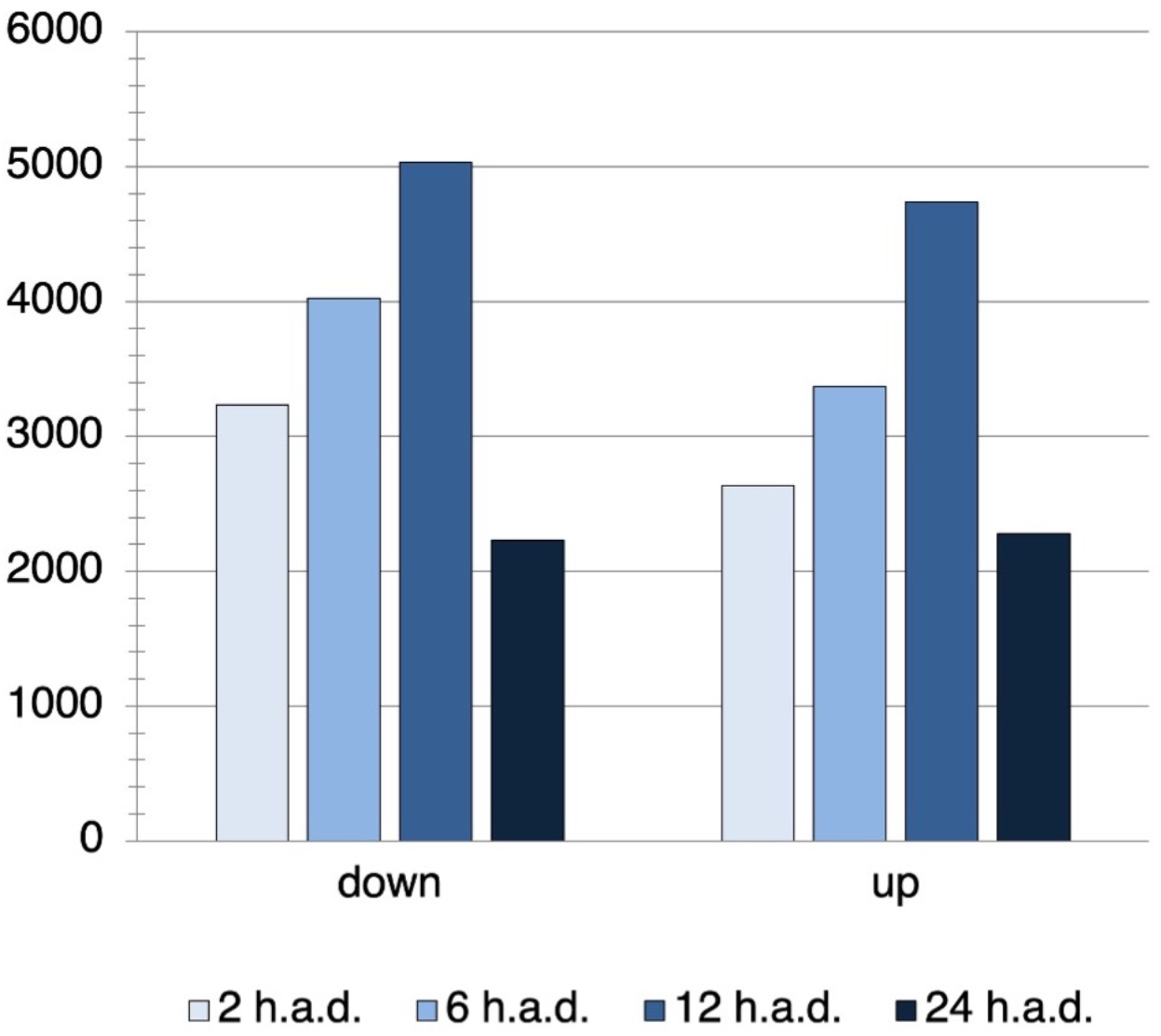
Number of genes repressed or induced 2, 6, 12 and 24 hours after decapitation.

**Figure S2.**
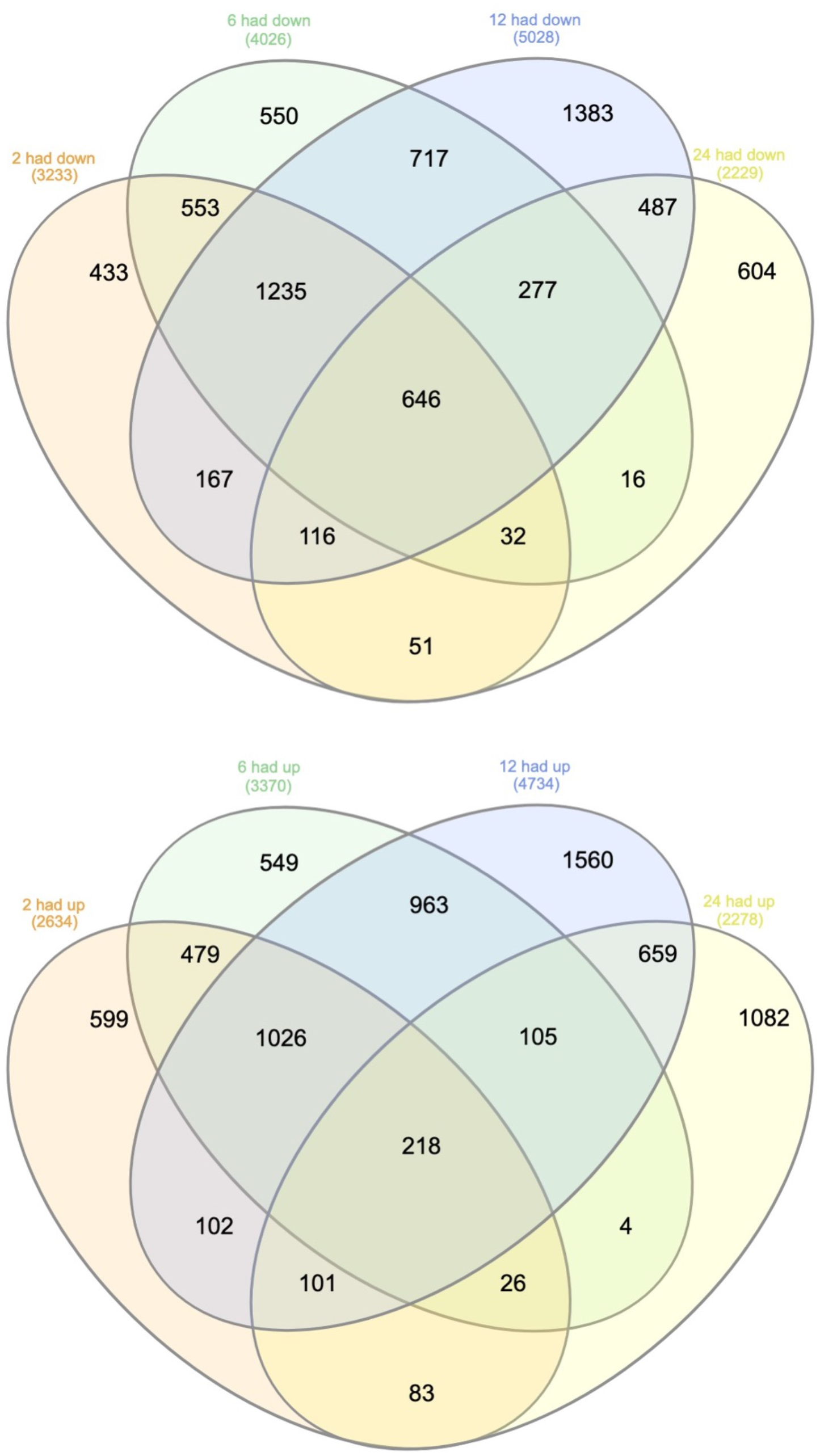
Venn diagrams for all genes repressed or induced 2, 6, 12 and 24 hours after decapitation.

**Figure S3.**
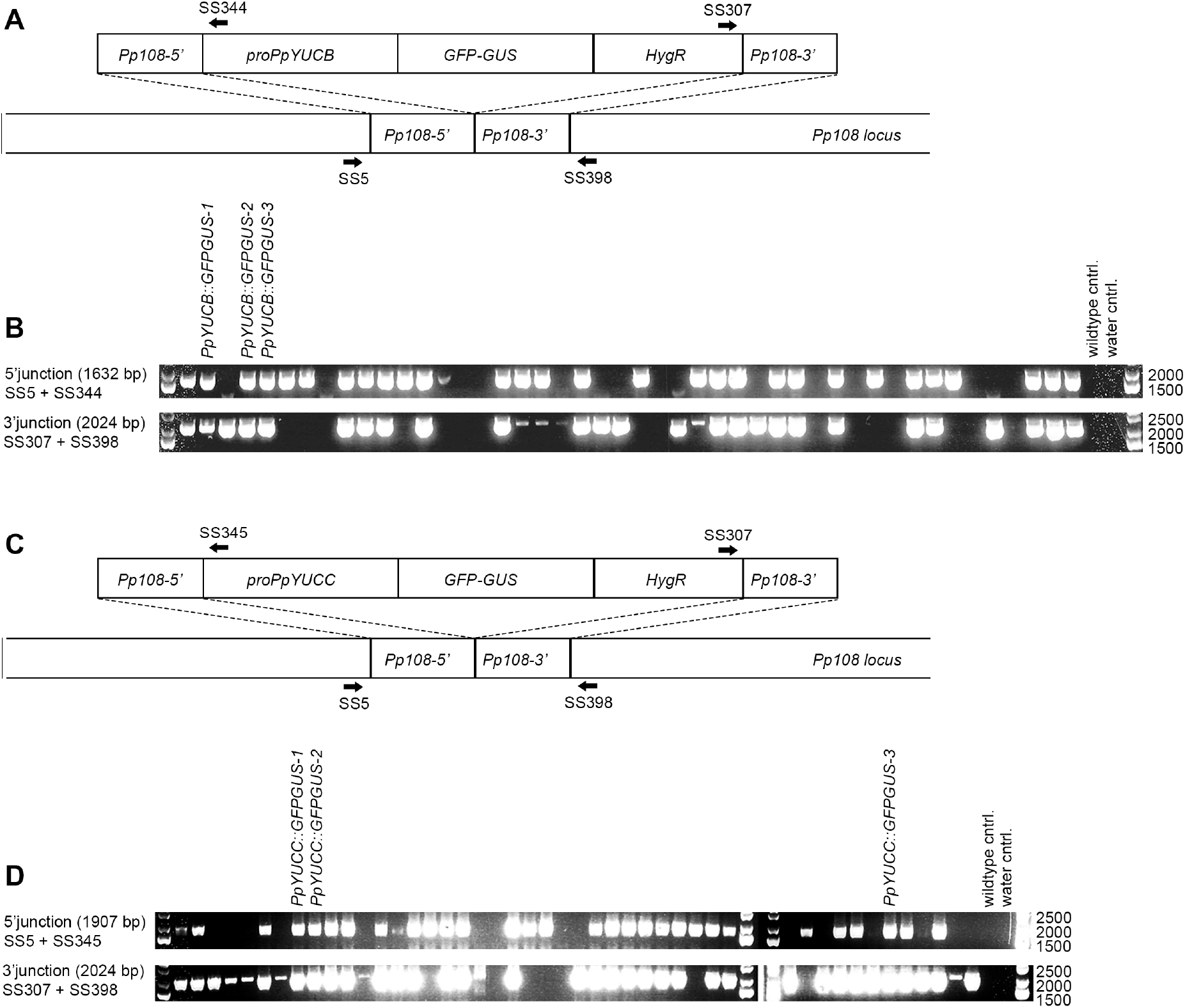
PCR verification of *PpYUCB and PpYUCC* transcriptional reporter lines in WT background. (**A**) Schematic view of the *PpYUCB::GFPGUS* reporter construct pMT244 and the *Pp108* locus to which it was targeted. Arrows mark the approximate annealing sites of primers used for PCR verification in B. (**B**) PCR verification of 5’ and 3’ junctions to confirm correct integration of the *PpYUCB::GFPGUS* reporter construct into the Pp108 locus. (**C**) Schematic view of the *PpYUCC::GFPGUS* reporter construct pMT251 and the *Pp108* locus to which it was targeted. Arrows mark the approximate annealing sites of primers used for PCR verification in D. (**D**) PCR verification of 5’ and 3’ junctions to confirm correct integration of the *PpYUCC::GFPGUS* reporter construct into the Pp108 locus. In both B and D, expected product sizes are indicated within parentheses. For primer sequences, see Table S5.

**Figure S4.**
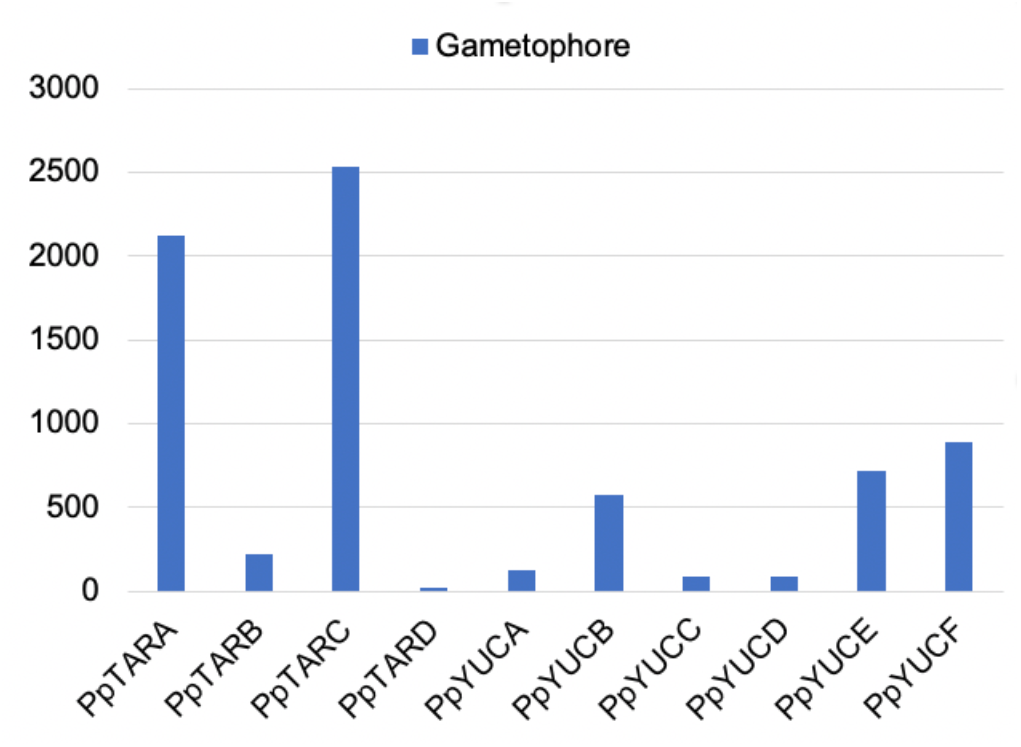
Absolute expression levels of *TAR* and *YUC* genes in *Physcomitrium patens* gametophores retrieved from Ortiz-Ramirez *et al*. (2016)^50^. Correspondence between gene names and identifiers: *PpTARA* (Pp3c21_15370V3.1, Pp1s167_103V6.1), *PpTARB* (Pp3c18_15140V3.1, Pp1s3_273V6.1), *PpTARC* (Pp3c17_6500V3.1, Pp1s26_28V6.1), *PpTARD* (Pp3c26_12520V3.1, Pp1s6_329V6.1), *PpYUCA* (Pp3c3_18590V1.1, Pp1s312_60V6.1), *PpYUCB* (Pp3c11_11790V3.1, Pp1s11_6V6.1), *PpYUCC* (Pp3c1_11500V3.1, Pp1s139_131V6.1), *PpYUCD* (Pp3c2_27740V3.1, Pp1s22_291V6.1), *PpYUCE* (Pp3c13_21970V3.1, Pp1s37_90V6.1), *PpYUCF* (Pp3c3_20490V3.1, Pp1s204_126V6.1).

**Figure S5.**
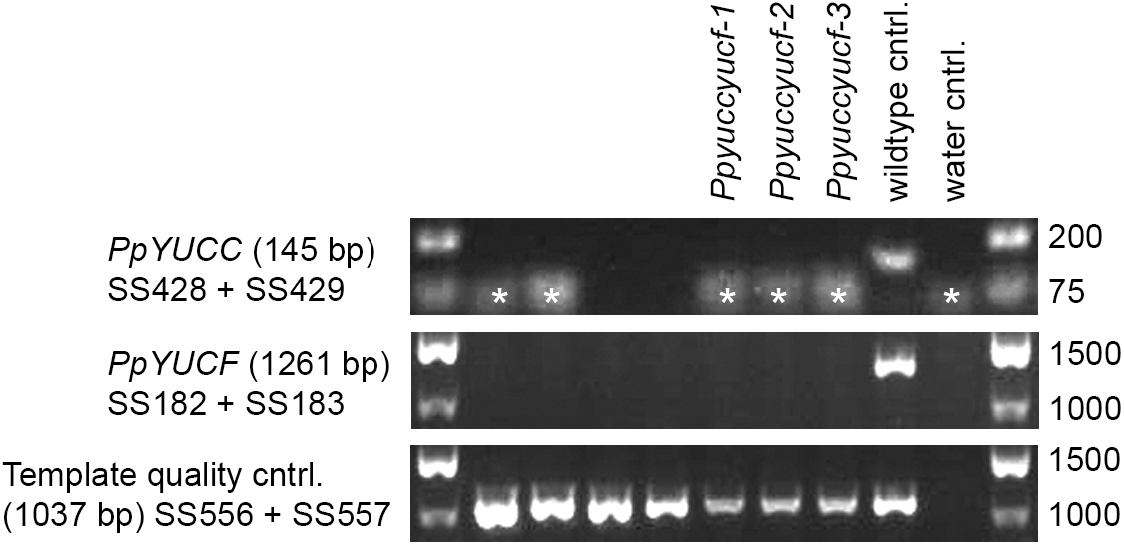
PCR genotyping of *yuccf* double knockout lines produced by a sexual cross of the single knockout lines *Ppyucc-2* and *Ppyucf-1*. (Landberg et al., 2020)^18^. The upper panel shows the result for a PCR confirming the loss of an internal *PpYUCC* gene sequence. Asterisks mark primer dimers. The middle panel shows the result for a PCR confirming the loss of an internal *PpYUCF* gene sequence. The bottom panel shows amplification of an unrelated locus to confirm integrity of the gDNA used as template in all three panels. Next to each panel, the primers used and the expected product size are noted. For primer sequences, see Table S5.

**Figure S6.**
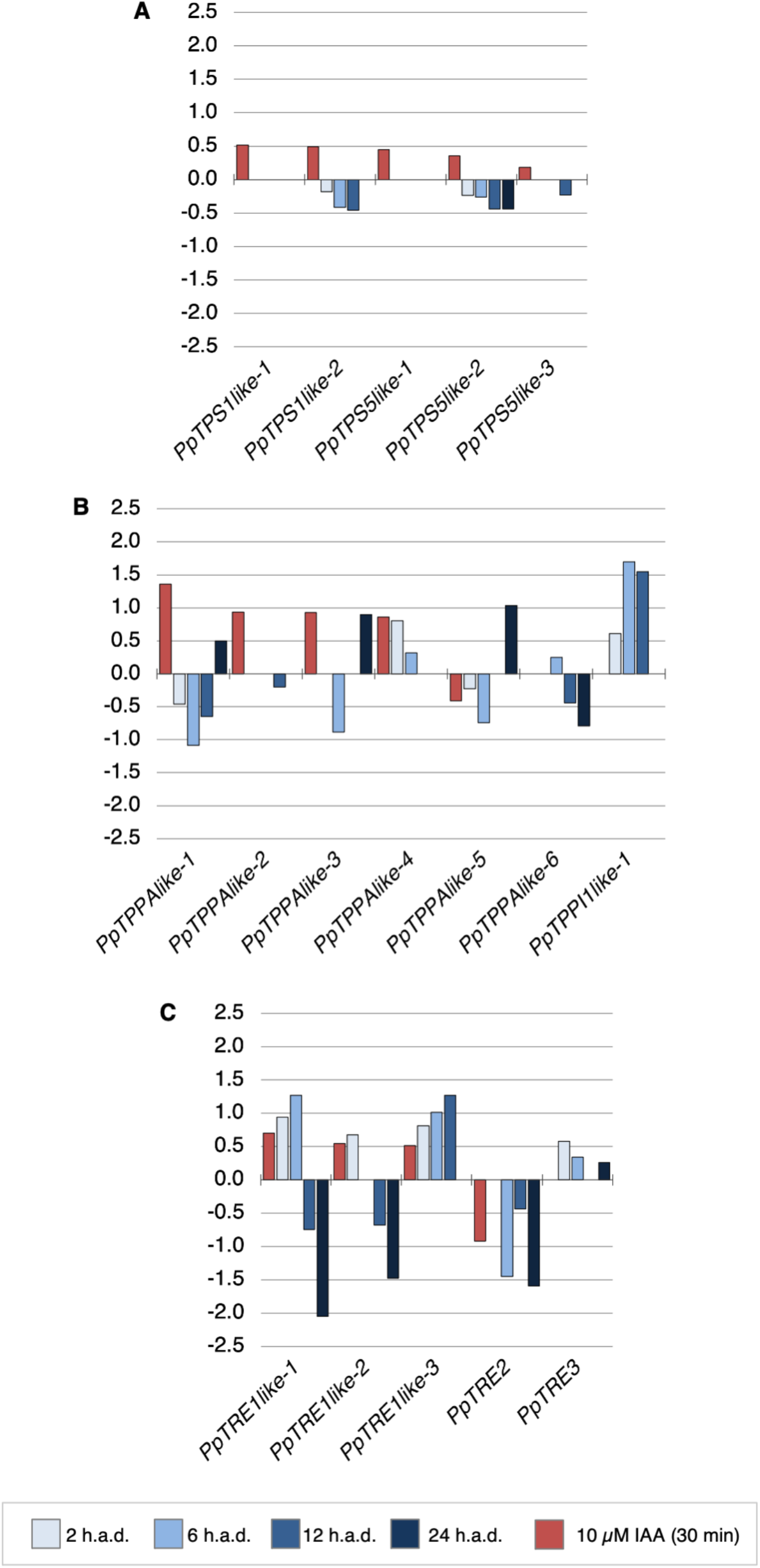
Expression of trehalose-6-phosphate (T6P) metabolism genes is affected by exogenous auxin and gametophore decapitation. Bar plots show *log2* fold-change in expression of *T6P synthase (PpTPS), T6P phosphatase (PpTPP)* and *trehalase (PpTRE)* coding genes in wild-type *Physcomitrium patens* gametophores after treatment with auxin (red bars) or decapitation (blue bars). Correspondence between gene names and identifiers: *PpTPPAlike-1* (Pp3c12_20050V3.1), *PpTPPAlike-2* (Pp3c4_20080V3.1), *PpTPPAlike-3* (Pp3c4_11080V3.1), *PpTPPAlike-4* (Pp3c10_13170V3.1), *PpTPPAlike-5* (Pp3c3_25080V3.1), *PpTPPAlike-6* (Pp3c3_22990V3.1), *PpTPPC1like-1* (Pp3c12_7510V3.1), *PpTPPE1like-1* (Pp3c12_7530V3.1), *PpTPPI1like-1* (Pp3c10_23130V3.1), *PpTPS1like-1* (Pp3c5_17730V3.1), *PpTPS1like-2* (Pp3c6_16450V3.1), *PpTPS5like-1* (Pp3c25_6990V3.1), *PpTPS5like-2* (Pp3c11_17560V3.1), *PpTPS5like-3* (Pp3c7_15250V3.1), *PpTRE1like-1* (Pp3c5_16470V3.1), *PpTRE1like-2* (Pp3c23_11240V3.1), *PpTRE1like-3* (Pp3c16_7830V3.1), *PpTRE1like-4* (Pp3c24_9748V3.1), *PpTRE1like-5* (Pp3c24_9750V3.1), *PpTRE2* (Pp3c6_4940V3.1), *PpTRE3* (Pp3c10_5310V3.1).

## Acknowledgements

YC acknowledges Fabrice Besnard, Joe Cammarata and Teva Vernoux for constructive discussions and suggestions on a draft version of this article, Stéphanie Hallet for technical support, and the CNRS (ATIP-Avenir programme) for research funding. The sequencing platform (POPS) benefited from the support of the LabEx Saclay Plant Sciences-SPS (ANR-10-LABX-0040-SPS).

## References

1. Coudert, Y., Bell, N.E., Edelin, C., and Harrison, C.J. (2017). Multiple innovations underpinned branching form diversification in mosses. New Phytol 22, 810.

2. Coudert, Y., Palubicki, W., Ljung, K., Novak, O., Leyser, O., and Harrison, C.J. (2015). Three ancient hormonal cues co-ordinate shoot branching in a moss. eLife 4, e06808.

3. Domagalska, M.A., and Leyser, O. (2011). Signal integration in the control of shoot branching. Nat Rev Mol Cell Biol 12, 211–221.

4. Ferguson, B.J., and Beveridge, C.A. (2009). Roles for auxin, cytokinin, and strigolactone in regulating shoot branching. Plant Physiol 149, 1929–1944.

5. Coudert, Y., Harris, S., and Charrier, B. (2019). Design Principles of Branching Morphogenesis in Filamentous Organisms. Curr. Biol. 29, R1149–R1162.

6. Barthélémy, D., and Caraglio, Y. (2007). Plant architecture: a dynamic, multilevel and comprehensive approach to plant form, structure and ontogeny. Ann Bot 99, 375–407.

7. Coudert, Y. (2017). The Evolution of Branching in Land Plants: Between Conservation and Diversity. In Evolutionary Developmental Biology (Springer International Publishing), pp. 1–17.

8. Cline, M. (1997). Concepts and terminology of apical dominance. Am J Bot 84, 1064.

9. Von Maltzahn, K. (1959). Interaction between Kinetin and Indoleacetic Acid in the Control of Bud Reactivation in Splachnum ampullaceum (L.) Hedw. Nature 183, 60–61.

10. Eklund, D.M., Thelander, M., Landberg, K., Ståldal, V., Nilsson, A., Johansson, M., Valsecchi, I., Pederson, E.R.A., Kowalczyk, M., Ljung, K., et al. (2010). Homologues of the Arabidopsis thaliana SHI/STY/LRP1 genes control auxin biosynthesis and affect growth and development in the moss Physcomitrella patens. Development 137, 1275–1284.

11. Sohlberg, J.J., Myrenås, M., Kuusk, S., Lagercrantz, U., Kowalczyk, M., Sandberg, G., and Sundberg, E. (2006). STY1 regulates auxin homeostasis and affects apical-basal patterning of the Arabidopsis gynoecium. The Plant Journal 47, 112–123.

12. Lavy, M., Prigge, M.J., Tao, S., Shain, S., Kuo, A., Kirchsteiger, K., Estelle, M., and Hardtke, C.S. (2016). Constitutive auxin response in Physcomitrella reveals complex interactions between Aux/IAA and ARF proteins. eLife Sciences 5, e13325.

13. Thelander, M., Landberg, K., and Sundberg, E. (2018). Auxin-mediated developmental control in the moss Physcomitrella patens. J Exp Bot 69, 277–290.

14. Yadav, S., Kumar, H., and Yadav, R.K. (2020). Local auxin biosynthesis promotes shoot patterning and stem cell differentiation in Arabidopsis shoot apex. bioRxiv, 819342.

15. Galvan-Ampudia, C.S., Cerutti, G., Legrand, J., Brunoud, G., Martin-Arevalillo, R., Azais, R., Bayle, V., Moussu, S., Wenzl, C., Jaillais, Y., et al. (2020). Temporal integration of auxin information for the regulation of patterning. eLife 9, e55832.

16. Cheng, Y., Dai, X., and Zhao, Y. (2006). Auxin biosynthesis by the YUCCA flavin monooxygenases controls the formation of floral organs and vascular tissues in Arabidopsis. Genes Dev. 20, 1790–1799.

17. Ljung, K. (2013). Auxin metabolism and homeostasis during plant development. Development 140, 943–950.

18. Landberg, K., Šimura, J., Ljung, K., Sundberg, E., and Thelander, M. (2021). Studies of moss reproductive development indicate that auxin biosynthesis in apical stem cells may constitute an ancestral function for focal growth control. New Phytologist 229, 845–860.

19. Kawai, Y., Ono, E., and Mizutani, M. (2014). Evolution and diversity of the 2-oxoglutarate-dependent dioxygenase superfamily in plants. Plant J 78, 328–343.

20. Muff, S., Nilsen, E.B., O’Hara, R.B., and Nater, C.R. (2021). Rewriting results sections in the language of evidence. Trends in Ecology& Evolution 0.

21. Bennett, T.A., Liu, M.M., Aoyama, T., Bierfreund, N.M., Braun, M., Coudert, Y., Dennis, R.J., O’Connor, D., Wang, X.Y., White, C.D., et al. (2014). Plasma Membrane-Targeted PIN Proteins Drive Shoot Development in a Moss. Curr. Biol. 24, 2776–2785.

22. Lang, D., Ullrich, K.K., Murat, F., Fuchs, J., Jenkins, J., Haas, F.B., Piednoel, M., Gundlach, H., Van Bel, M., Meyberg, R., et al. (2018). The Physcomitrella patens chromosome-scale assembly reveals moss genome structure and evolution. The Plant Journal 93, 515–533.

23. Romani, F. (2017). Origin of TAA Genes in Charophytes: New Insights into the Controversy over the Origin of Auxin Biosynthesis. Front. Plant Sci. 8, R899–3.

24. Delaux, P.-M., Hetherington, A.J., Coudert, Y., Delwiche, C., Dunand, C., Gould, S., Kenrick, P., Li, F.-W., Philippe, H., Rensing, S.A., et al. (2019). Reconstructing trait evolution in plant evo-devo studies. Curr. Biol. 29, R1110–R1118.

25. Eklund, D.M., Ishizaki, K., Flores-Sandoval, E., Kikuchi, S., Takebayashi, Y., Tsukamoto, S., Hirakawa, Y., Nonomura, M., Kato, H., Kouno, M., et al. (2015). Auxin Produced by the Indole-3-Pyruvic Acid Pathway Regulates Development and Gemmae Dormancy in the Liverwort Marchantia polymorpha. The Plant Cell Online 27, 1650–1669.

26. Solly, J.E., Cunniffe, N.J., and Harrison, C.J. (2017). Regional Growth Rate Differences Specified by Apical Notch Activities Regulate Liverwort Thallus Shape. Current Biology 27, 16–26.

27. Shubin, N., Tabin, C., and Carroll, S. (2009). Deep homology and the origins of evolutionary novelty. Nature 457, 818–823.

28. Véron, E., Vernoux, T., and Coudert, Y. (2020). Phyllotaxis from a Single Apical Cell. Trends in Plant Science.

29. Toyokura, K., Goh, T., Shinohara, H., Shinoda, A., Kondo, Y., Okamoto, Y., Uehara, T., Fujimoto, K., Okushima, Y., Ikeyama, Y., et al. (2019). Lateral Inhibition by a Peptide Hormone-Receptor Cascade during Arabidopsis Lateral Root Founder Cell Formation. Dev Cell 48, 64–75.e5.

30. Chandler, J.W., and Werr, W. (2017). DORNRÖSCHEN, DORNRÖSCHEN-LIKE, and PUCHI redundantly control floral meristem identity and organ initiation in Arabidopsis. J Exp Bot 68, 3457–3472.

31. Capua, Y., and Eshed, Y. (2017). Coordination of auxin-triggered leaf initiation by tomato LEAFLESS. Proc Natl Acad Sci USA 114, 3246–3251.

32. Ishikawa, M., Morishita, M., Higuchi, Y., Ichikawa, S., Ishikawa, T., Nishiyama, T., Kabeya, Y., Hiwatashi, Y., Kurata, T., Kubo, M., et al. (2019). Physcomitrella STEMIN transcription factor induces stem cell formation with epigenetic reprogramming. Nat Plants 5, 681–690.

33. Fichtner, F., Barbier, F.F., Feil, R., Watanabe, M., Annunziata, M.G., Chabikwa, T.G., Höfgen, R., Stitt, M., Beveridge, C.A., and Lunn, J.E. (2017). Trehalose 6-phosphate is involved in triggering axillary bud outgrowth in garden pea (Pisum sativum L.). The Plant Journal 92, 611–623.

34. Fichtner, F., Barbier, F.F., Annunziata, M.G., Feil, R., Olas, J.J., Mueller-Roeber, B., Stitt, M., Beveridge, C.A., and Lunn, J.E. (2021). Regulation of shoot branching in arabidopsis by trehalose 6-phosphate. New Phytol 229, 2135–2151.

35. Satoh-Nagasawa, N., Nagasawa, N., Malcomber, S., Sakai, H., and Jackson, D. (2006). A trehalose metabolic enzyme controls inflorescence architecture in maize. Nature 441, 227–230.

36. Claeys, H., Vi, S.L., Xu, X., Satoh-Nagasawa, N., Eveland, A.L., Goldshmidt, A., Feil, R., Beggs, G.A., Sakai, H., Brennan, R.G., et al. (2019). Control of meristem determinacy by trehalose 6-phosphate phosphatases is uncoupled from enzymatic activity. Nature Plants, 1–9.

37. Hiss, M., Meyberg, R., Westermann, J., Haas, F.B., Schneider, L., Schallenberg-Rüdinger, M., Ullrich, K.K., and Rensing, S.A. (2017). Sexual reproduction, sporophyte development and molecular variation in the model moss Physcomitrella patens: introducing the ecotype Reute. The Plant Journal 22, 9.

38. Thelander, M., Landberg, K., and Sundberg, E. (2019). Minimal auxin sensing levels in vegetative moss stem cells revealed by a ratiometric reporter. New Phytologist 224, 775–788.

39. Schaefer, D., Zryd, J.P., Knight, C.D., and Cove, D.J. (1991). Stable transformation of the moss Physcomitrella patens. Mol Gen Genet 226, 418–424.

40. Rensing, S.A., Lang, D., Zimmer, A.D., Terry, A., Salamov, A., Shapiro, H., Nishiyama, T., Perroud, P.-F., Lindquist, E.A., Kamisugi, Y., et al. (2008). The Physcomitrella genome reveals evolutionary insights into the conquest of land by plants. Science 319, 64–69.

41. Gagnot, S., Tamby, J.-P., Martin-Magniette, M.-L., Bitton, F., Taconnat, L., Balzergue, S., Aubourg, S., Renou, J.-P., Lecharny, A., and Brunaud, V. (2008). CATdb: a public access to Arabidopsis transcriptome data from the URGV-CATMA platform. Nucleic Acids Research 36, D986–D990.

42. Kopylova, E., Noé, L., and Touzet, H. (2012). SortMeRNA: fast and accurate filtering of ribosomal RNAs in metatranscriptomic data. Bioinformatics 28, 3211–3217.

43. Langmead, B., and Salzberg, S.L. (2012). Fast gapped-read alignment with Bowtie 2. Nat Methods 9, 357–359.

44. Rigaill, G., Balzergue, S., Brunaud, V., Blondet, E., Rau, A., Rogier, O., Caius, J., Maugis-Rabusseau, C., Soubigou-Taconnat, L., Aubourg, S., et al. (2018). Synthetic data sets for the identification of key ingredients for RNA-seq differential analysis. Briefings in Bioinformatics 19, 65–76.

45. McCarthy, D.J., Chen, Y., and Smyth, G.K. (2012). Differential expression analysis of multifactor RNA-Seq experiments with respect to biological variation. Nucleic Acids Research 40, 4288–4297.

46. Heberle, H., Meirelles, G.V., da Silva, F.R., Telles, G.P., and Minghim, R. (2015). InteractiVenn: a web-based tool for the analysis of sets through Venn diagrams. BMC Bioinformatics 16, 169.

47. Ge, S.X., Jung, D., and Yao, R. (2020). ShinyGO: a graphical gene-set enrichment tool for animals and plants. Bioinformatics 36, 2628–2629.

48. Edgar, R., Domrachev, M., and Lash, A.E. (2002). Gene Expression Omnibus: NCBI gene expression and hybridization array data repository. Nucleic Acids Research 30, 207–210.

49. Kleiber, C., and Zeileis, A. (2020). AER: Applied Econometrics with R.

50. Ortiz-Ramírez, C., Hernandez-Coronado, M., Thamm, A., Catarino, B., Wang, M., Dolan, L., Feijó, J.A., and Becker, J.D. (2016). A Transcriptome Atlas of Physcomitrella patens Provides Insights into the Evolution and Development of Land Plants. Mol Plant 9, 205–220.

